# m6A methylation potentiates cytosolic dsDNA recognition in a sequence-specific manner

**DOI:** 10.1101/742072

**Authors:** Melania Balzarolo, Sander Engels, Anja J. de Jong, Katka Franke, Timo K. van den Berg, Edith M. Janssen, Bas van Steensel, Monika C. Wolkers

**Affiliations:** Sanquin Research, Department of Hematopoiesis, and Landsteiner Laboratory, Academic Medical Centre (AMC), University of Amsterdam, Amsterdam, The Netherlands; Sanquin Research, Department of Blood cell research, and Landsteiner Laboratory, Academic Medical Centre (AMC), University of Amsterdam, Amsterdam, The Netherlands; Division of Molecular Immunology, Cincinnati Children’s Hospital Research Foundation, University of Cincinnati College of Medicine, Room S5.419, 3333 Burnet Avenue, Cincinnati, OH 45229; Division of Gene Regulation, Netherlands Cancer Institute, 1066 CX Amsterdam, Netherlands

## Abstract

Nucleic acid sensing through pattern recognition receptors is critical for immune recognition of microbial infections. Microbial DNA is frequently methylated at the N^6^ position of adenines (m6A), a modification that is rare in mammalian host DNA. We show that m6A-methylation of 5’-GATC-3’ motifs augments the immunogenicity of double stranded (ds)DNA in macrophages and dendritic cells. Transfection with m6A-methylated DNA increased the expression of the activation markers CD69 and CD86, and of *Ifn*β, *iNos* and *Cxcl10* mRNA. Recognition of m6A DNA occurs independently of TLR and RIG-I signaling but requires STING, a key mediator of cytosolic DNA sensing. Intriguingly, the response to m6A DNA is sequence-specific. m6A is immunostimulatory in some motifs, but immunosuppressive in others, a feature that is conserved between mouse and human. In conclusion, epigenetic alterations of bacterial DNA are differentially perceived by innate cells, a feature that could potentially be used for the design of immune-modulating therapeutics.

## Introduction

Innate immune cells can recognize invading pathogens through pattern recognition receptors (PRRs) (1). This feature allows for rapid recognition of invading pathogens and for a swift onset of immune responses. De-regulation of PRR sensing signaling is associated with pathogenic and autoimmune conditions (2, 3).

A wide range of PRRs localize in the endosomes and in the cytosol where they detect bacterial and viral nucleic acids (3–5). In the endosome, Toll-like receptors (TLRs) sense single-stranded (ss) and double-stranded (ds)RNA (TLR3), as well as conserved pathogen-derived ssDNA structures (TLR7/9) (6). Engaging these TLRs leads to the induction of pro-inflammatory cytokines like Interleukin (IL)-6, Tumor necrosis factor (TNF)-α, and type I Interferons (IFNs) in an NF-kB- and MYD88/TRIF-dependent manner (6–9). In the cytosol, viral dsRNA is recognized by the RIG-I-like family of receptors (RLRs) and MDA5 (5). Through the adaptor protein IPS1/MAVS, proinflammatory cytokines and type I IFNs are produced (5, 10). dsDNA present in the cytosol is primarily recognized by cGAS and AIM2, which promote the production of type I IFNs and IL-1β through STING and ASC, respectively (11, 12). Other DNA sensors include RNA polymerase III, IFI16 and DAI (4, 5).

Recognition of pathogenic cytosolic DNA is influenced by sequence length, secondary structures and nucleotide overhangs (3, 5). For instance, the right-handed (B) form of DNA is well recognized by cytosolic DNA sensors (13, 14). Furthermore, guanosine overhangs in conserved Y-form DNA of retroviruses such as the human immunodeficiency virus type 1 (HIV-1) potentiate type I IFN production in human macrophages (15).

Eukaryotic and microbial DNA also differ in their epigenetic landscape, in particular methylation of adenines and cytosines. These modifications are catalyzed by DNA methyltransferases (MTases). Adenine and cytosine methylations are found in DNA of most prokaryotes (16) and are involved in bacterial defense, virulence, chromosomal replication, and gene regulation (16, 17). The best studied prokaryotic MTase is DNA adenine methyltransferase (Dam). Dam was originally described in *Escherichia coli* and methylates adenine in position N^6^ (m6A) in 5’-GATC-3’ DNA motifs, generating a G^m6^ATC DNA motif (18). Other sequence motifs in a variety of prokaryotes can also carry m6A, depending on the species (16).

Potentially, differences in methylation status could be used by the innate immune system to discriminate pathogen-derived DNA from host DNA. For example, CpG motifs are mostly unmethylated in microbial genomes (16), but frequently methylated in DNA across a variety of human and mouse tissues (19, 20). This difference is recognized by the PRR TLR9 (16, 17), leading to the production of inflammatory cytokines. Thus, recognition of CpG motifs forms a prime example for immune cells to discriminate host DNA from the microbial genome.

Much less is known about a putative immunogenic role of m6A in DNA. This modification is present in human and mouse DNA, but it appears to be extremely rare (in the range of 0.0005 - 0.05% of all adenines) (21, 22) compared to the pervasive presence in prokaryotic DNA (16). This could thus be another basis for discrimination of host and pathogen DNA. Indeed, a previous study showed that systemic injection of DNA containing one G^m6^ATC motif resulted in increased blood levels on the proinflammatory cytokines TNF-α, IL-6 and IL-12 in mice (23). However, which cells respond to m6A-methylated DNA and through which innate immune sensors has not been studied (24). Furthermore, it is not known whether m6A recognition is restricted to G^m6^ATC motifs or whether it is also observed in another sequence context.

Here, we interrogated whether cytosolic delivery of G^m6^ATC DNA provokes immune cell response in innate immune cells, and if so, through which mechanism. We found that synthetic dsDNA containing G^m6^ATC motifs potentiates the response of murine macrophages and dendritic cells. This recognition requires STING-mediated signaling. Importantly, m6A-methylation does not boost immune responses *per se*, but depends on the nucleotide sequence context, a feature that is conserved in mouse and in human macrophages.

## Materials and Methods

### Mice

C57BL/6J mice (bred at the animal department of the Netherlands Cancer Institute, Amsterdam, The Netherlands), or mice deficient for MYD88xTRIF ((8, 25) hereafter *Myd88*^-/-^ *Trif*^-/-^), for IPS-1 ((26), *Ips*^-/-^), or for STING ((27), *Sting*^-/-^) were used. All animal experiments were performed in accordance with institutional and national guidelines and approved by the Experimental Animal Committee of the Netherlands Cancer Institute, and of the Cincinnati Children’s Hospital.

### Generation of murine bone marrow-derived macrophages and dendritic cells

Bone marrow (BM) cells were obtained from mouse tibias and femurs. Briefly, after BM was flushed from the bones, red blood cells were lysed with red blood cell lysis buffer containing 0.168 M NH_4_Cl, and washed once with PBS (28). Bone marrow-derived macrophages (BMMs) were generated by seeding 2 x 10^6^ BM cells in a 100 mm non-tissue culture treated dish in RPMI 1640 (Lonza) supplemented with 10% FCS, 2 mM L-glutamine, 100 U/mL penicillin, 100 μg/mL streptomycin and β-mercaptoethanol together with 15% L-929 conditioned medium containing recombinant M-CSF for 8 days at 37°C and 5% CO_2_. Medium was refreshed after 4 days.

Bone marrow-derived dendritic cells were generated with recombinant Flt3L (Flt3L-DCs) as previously described (28). Briefly, BM cells were cultured at 1.5 × 10^6^ cells/ml for 9-10 days at 37°C and 5% CO_2_ in complete DC medium (RPMI 1640 supplemented with 5% FCS, 2 mM L-glutamine, 100 U/mL penicillin, 100 μg/mL streptomycin, and β-mercaptoethanol) supplemented with 30% conditioned medium from CHO cells producing murine recombinant Flt3L (29). BMMs and Flt3L-DC cultures were 95-99% F4/80^+^ or CD11c^+^, respectively.

### Generation of human monocyte-derived macrophages

Peripheral mononuclear blood cells (PBMC) were isolated from peripheral blood or buffy coats of healthy individuals collected by Sanquin Blood Supply (Amsterdam, Netherlands). The study was performed according to the Declaration of Helsinki (seventh revision, 2013). Written informed consent was obtained (Sanquin, Amsterdam, The Netherlands). Monocyte isolation was performed by gradient centrifugation on Percoll (Pharmacia, Uppsala, Sweden) following by magnetic-activated cell separation sorting using human CD14 Microbeads (Miltenyi Biotec). Freshly isolated CD14^+^ monocytes were cultured for 7-8 days to differentiate into macrophages in IMDM medium supplemented with 10% FCS, 100 U/ml penicillin, 100 µg/ml streptomycin, 2 mM L-glutamine and 20 ng/ml human macrophage colony-stimulating factor (M-CSF) (eBioscience).

### Generation of double stranded GATC and G^m6^ATC sequences

HPLC-grade DNA oligos (Sigma-Aldrich) were dissolved in sterile endotoxin-free water, aliquoted and stored at −20°C. To generate dsDNA, equimolar amounts of m6A-methylated or unmethylated complementary oligos were linearized at 95°C, annealed at 75°C for 5 minutes, and slowly cooled down to room temperature. Double stranded sequences were aliquoted and stored at −20°C. dsDNA of GATC DNA was generated from multiple batches. For Tm analysis of each batch, 1 μg dsDNA was incubated with Sybr Green mix (Applied Biosystems) for 5 min at room temperature. Melting curve was determined on the Step-OnePlus™ Real-Time PCR System (Applied Biosystems) with the standard temperature gradient from 40-95°C.

### Stimulation and nucleic acid transfection

After generation, murine BMMs and Flt3L-DCs, and human monocyte-derived macrophages were seeded for 1 h at 37°C and 5% CO_2_ in 24- or 48-well non-tissue culture treated plates (BD) at a density of 1-2 x 10^5^ cells/ml, before being cultured for indicated time points in FCS-free medium containing 1 µg/ml LPS (Invivogen), 1 μg/ml synthetic (B) form DNA analog poly(deoxyadenylic-deoxythymidylic) acid (poly(dA:dT)) (Invivogen) or 400 nM dsDNA containing GATC or G^m6^ATC sequences, or variants thereof. Cells were transfected with poly(dA:dT), m6A methylated or unmethylated dsDNA with 0.1% Lipofectamine 2000 (Invitrogen) according to the manufacturer’s protocol. Cells in medium alone (untransfected, ctrl) or in medium containing Lipofectamine 2000 (mock) were as controls for DNA stimulation and DNA transfection, respectively. After indicated time points, cells were harvested by scraping from culture plates for analysis.

### Antibodies and Flow cytometry

BMMs and Flt3L-DCs were stained with antibodies directed against murine F4/80-APC (clone BM8), CD69-FITC (clone H1.2F3), CD11c-APC (clone N418), and CD86-FITC (clone GL1) (eBioscience). Stainings were performed in the presence of anti-CD16/CD32 block (2.4G2; kind gift from Louis Boon, Bioceros). Flow cytometry was performed with LSRII (BD Biosciences), and data were analysed with FlowJo software v7.6.5. (Tree Star, Inc).

### Quantitative Reverse Transcriptase-PCR

Total RNA was extracted using TRIzol reagent (Invitrogen). cDNA was generated with SuperScript III reverse transcriptase (Invitrogen), dNTPs (Fermentas) and Random Primer (Promega) according to manufacturer’s protocol. Quantitative Reverse Transcriptase-PCR (RT-qPCR) was performed using SYBR Green mix on the Step-OnePlus™ System (Applied Biosystems). Primers used for gene expression analysis (Table 1) were validated by serial dilutions. Gene expression was normalized to *L32* (mouse genes) *or 18s* (human genes).

**Table 1:**
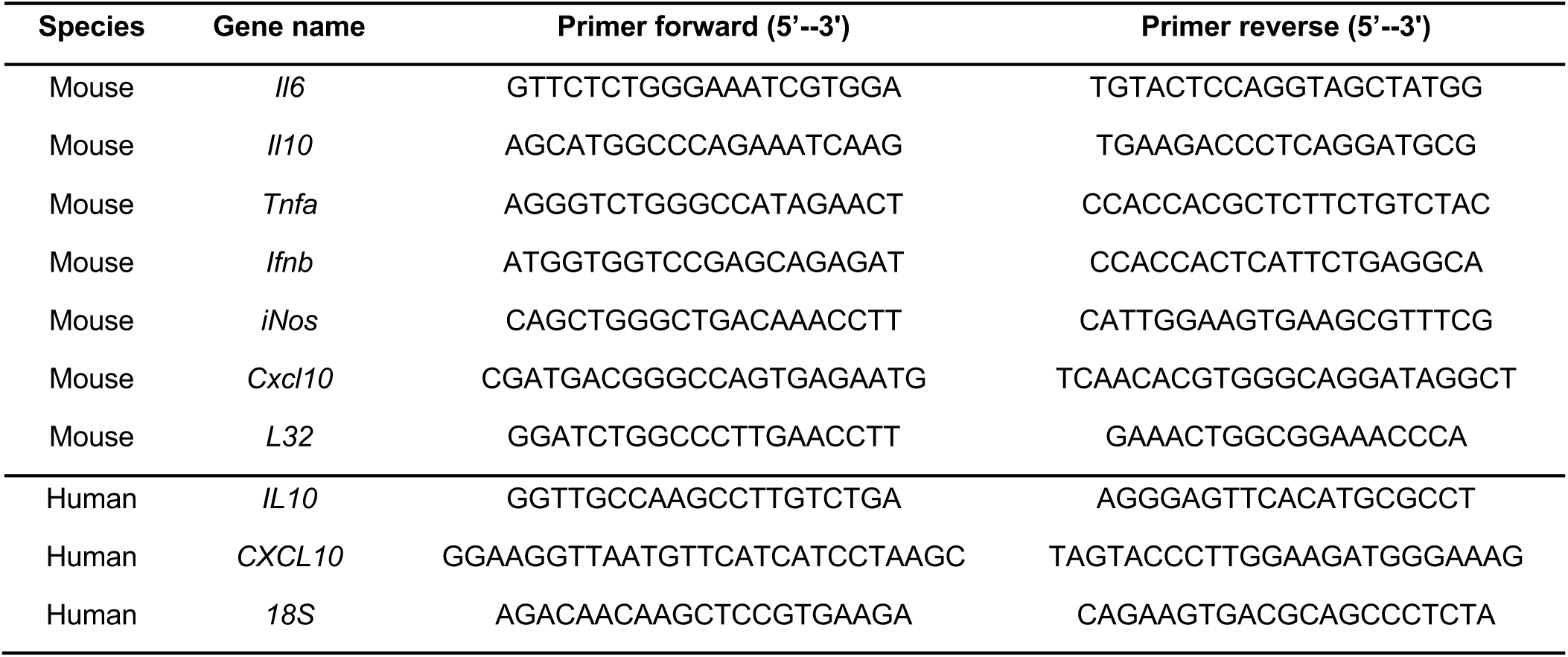
Primers used for RT-qPCR analysis.

### Statistical analysis

Data were analyzed for statistical significance with 2-tailed unpaired or paired Student’s *t*-test, as indicated (Prism v5, GraphPad Software). Results are expressed as mean ± standard deviation (SD) and were considered statistically significant with *p* values < 0.05.

## Results

### Cytosolic delivery of m6A-methylated dsDNA enhances macrophage and DC activation

We first examined whether N^6^-methyl-adenine (m6A) modifications in GATC motifs alters the immunogenicity of dsDNA for macrophages and dendritic cells. The sequence we selected for analysis is present in the genome of several bacterial strains, such as *Escherichia coli, Salmonella enterica* and *Klebsiella pneumoniae*. The 34bp long sequence contains a cluster of three GATC motifs but lacks CpG motifs (Table 2). To exclude other immune stimulants in the preparations, we used HPLC-purified oligos that were dissolved in endotoxin-free H2O. m6A modifications are abundant in bacteria on both DNA strands, which prompted us to study the response to double stranded DNA (dsDNA). We determined the integrity of the generated dsDNA by measuring the melting temperature (Tm) of the m6A-methylated (GATC DNA) or unmethylated (G^m6^ATC DNA) dsDNA. As expected, m6A modifications reduced the Tm of the dsDNA by ∼5°C, as a consequence of altering the struture and by destabilizing double stranded bonds (Table 2).

**Table 2:**
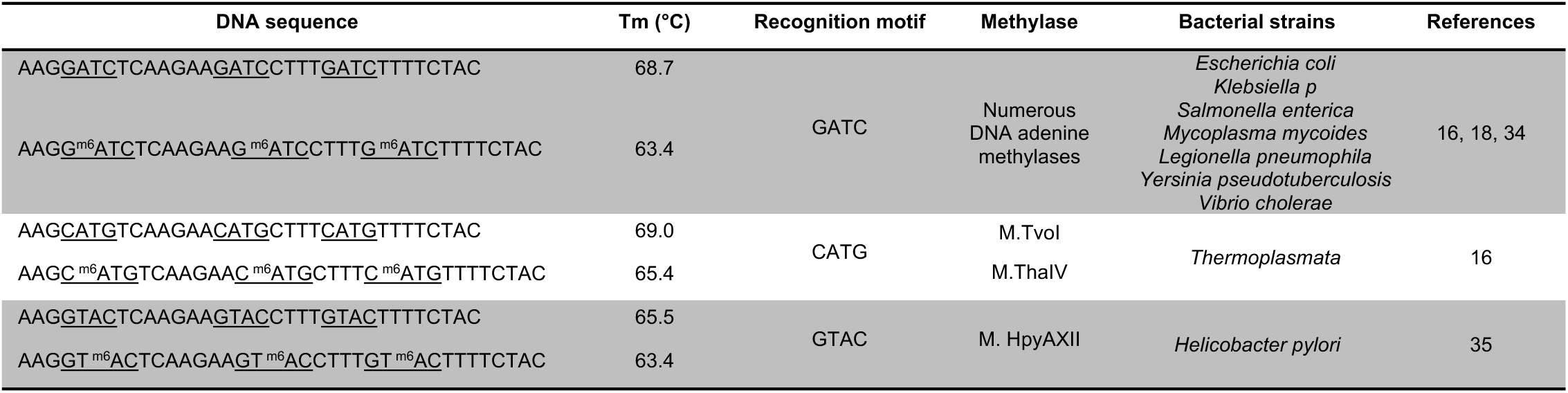
oligos and melting melting temperature (Tm) of corresponding dsDNA used in this study. Depicted are also the motifs recognized by prokaryotic methyltransferases (MTses), and examples of ial strains expressing the MTses.

Recognition of dsDNA by PRRs occurs primarily in the cytosol (3, 4). Therefore, to determine whether m6A modifications alter the immunogenicity of dsDNA, we delivered the dsDNA to marrow-derived macrophages from C57Bl/6J mice (BMMs) through transfection with Lipofectamine 2000. As a control, we transfected Poly(dA:dT), a well-studied (B) form dsDNA that elicits potent type I IFN response in both mouse and human cells (4). Within 6 h of stimulation BMMs transfected with poly(dA:dT) showed increased expression of CD69 (Fig. 1A), an early macrophage activation marker (8, 30). Transfection with the 34bp synthetic DNA sequences also resulted in increased CD69 expression (Fig. 1A). CD69 protein expression was even higher when cells were transfected G^m6^ATC DNA compared to unmethylated DNA (Fig. 1A). CD69 expression was also increased at later time points, i.e. 24 h after transfection with G^m6^ATC DNA (Fig. 1B). The induction of CD69 expression depended on intracellular delivery of the dsDNA, because the delivery of GATC or G^m6^ATC DNA without Lipofectamine 2000 did not induce expression of CD69 (Fig. 1B).

**Figure 1.**
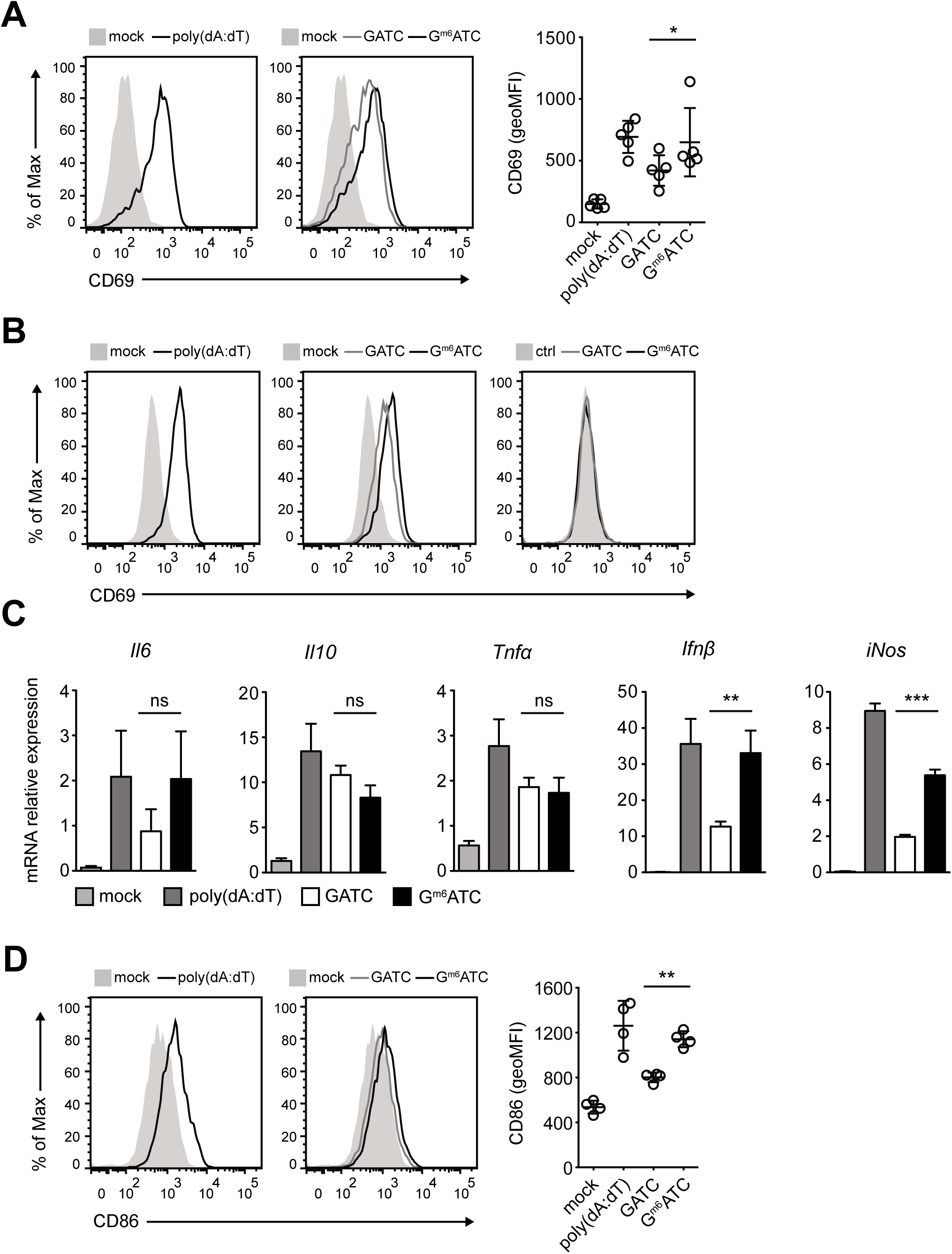
Cytosolic recognition of m6A-methylated dsDNA potentiates macrophage and dendritic cell activation. (**A**) Representative histogram of CD69 expression of bone marrow-derived macrophages (BMMs) 6 h after transfection with 0.1% Lipofectemine 2000 and 1 μg/ml poly(dA:dT) (*left panel*), 400 nM unmethylated (GATC) or 400 nM methylated (G^m6^ATC) DNA (*middle panel*). Transfection with 0.1% Lipofectamine 2000 alone served as control (mock). *Right panel:* CD69 expression levels (Geometric mean fluorescence intensity, geoMFI) compiled from five independently performed experiments. (**B**) CD69 expression of BMMs stimulated for 24 h with 1μg/ml poly(dA:dT), or with GATC or G^m6^ATC DNA in the presence (*middle panel*) or absence (*right panel*) of Lipofectemine. Lipofectamine mock treated or untreated BMMs (ctrl) served as controls. (**C**) *Il6, Il10, Tnfα, Ifnβ, and iNos* mRNA levels of BMMs activated for 6 h with indicated reagents. **B** and **C** are representative of two independently performed experiments. (**D**) Representative histograms (left) of CD86 expression and compiled data from 2 independently performed experiments (right) of BM-derived dendritic cells (Flt3L-DCs) that were mock transfected or transfected overnight with poly(dA:dT), GATC or G^m6^ATC DNA. Paired (A-E) or unpaired (C) Student’s *t*-test. (* *p* < 0.05, * * *p* < 0.01, * * * *p* < 0.001).

Macrophage activation with dsDNA leads to rapid transcription of inflammatory molecules (31). To determine whether m6A-methylation alters the inflammatory gene expression profile of macrophages, we measured the mRNA levels of *Il6, Il10, Tnfα, Ifnβ* and *iNos*. *Il6, Il10, and Tnfα* mRNA levels were increased upon transfection with both DNA variants, and it occurred irrespective of the methylation status of the dsDNA (Fig. 1C). We also observed increased mRNA levels of the early inflammatory genes *Ifnβ* and *iNos*, and both transcripts were more potently induced upon transfection with G^m6^ATC DNA (Fig. 1C; p=0.005 and p<0.0001, respectively). Similarly, bone-marrow derived DCs generated with Flt3L showed increased levels of the costimulatory molecule CD86 upon transfection with G^m6^ATC DNA when compared to transfection with GATC DNA (Fig. 1D). Thus, m6A modification in GATC motifs promotes the gene expression of several key inflammatory molecules.

### STING drives immune activation for both m6A-modified and unmodified DNA

We next interrogated which PRR mediates the recognition of the m6A-methylated dsDNA. TLR3, TLR7/8 and TLR9 which detect nucleic acids (32) signal through MYD88 and TRIF, the key adaptor molecules downstream of TLR signaling (8, 9). To determine whether TLRs can sense methylated dsDNA, we generated BMMs from *Myd88*^-/-^*Trif*^-/-^ mice. As expected, *Myd88*^-/-^*Trif*^-/-^ BMMs failed to respond to the TLR4 ligand LPS after 6 h of stimulation, but maintained their ability to respond to poly(dA:dT), which is sensed in an TLR-independent manner (14)(Fig. 2A, B). Transfection with GATC and G^m6^ATC DNA resulted in identical effects in *Myd88*^-/-^*Trif*^-/-^ and *wt* BMMs, with higher CD69 expression upon transfection with G^m6^ATC DNA (Fig. 2A, B). This suggests that TLRs are dispensable for dsDNA recognition. The adaptor protein IPS-1 that acts downstream of the dsRNA recognizing RIG-I-like receptors (26, 33) was also not required for either GATC, or G^m6^ATC DNA recognition (Fig. 2C).

**Figure 2.**
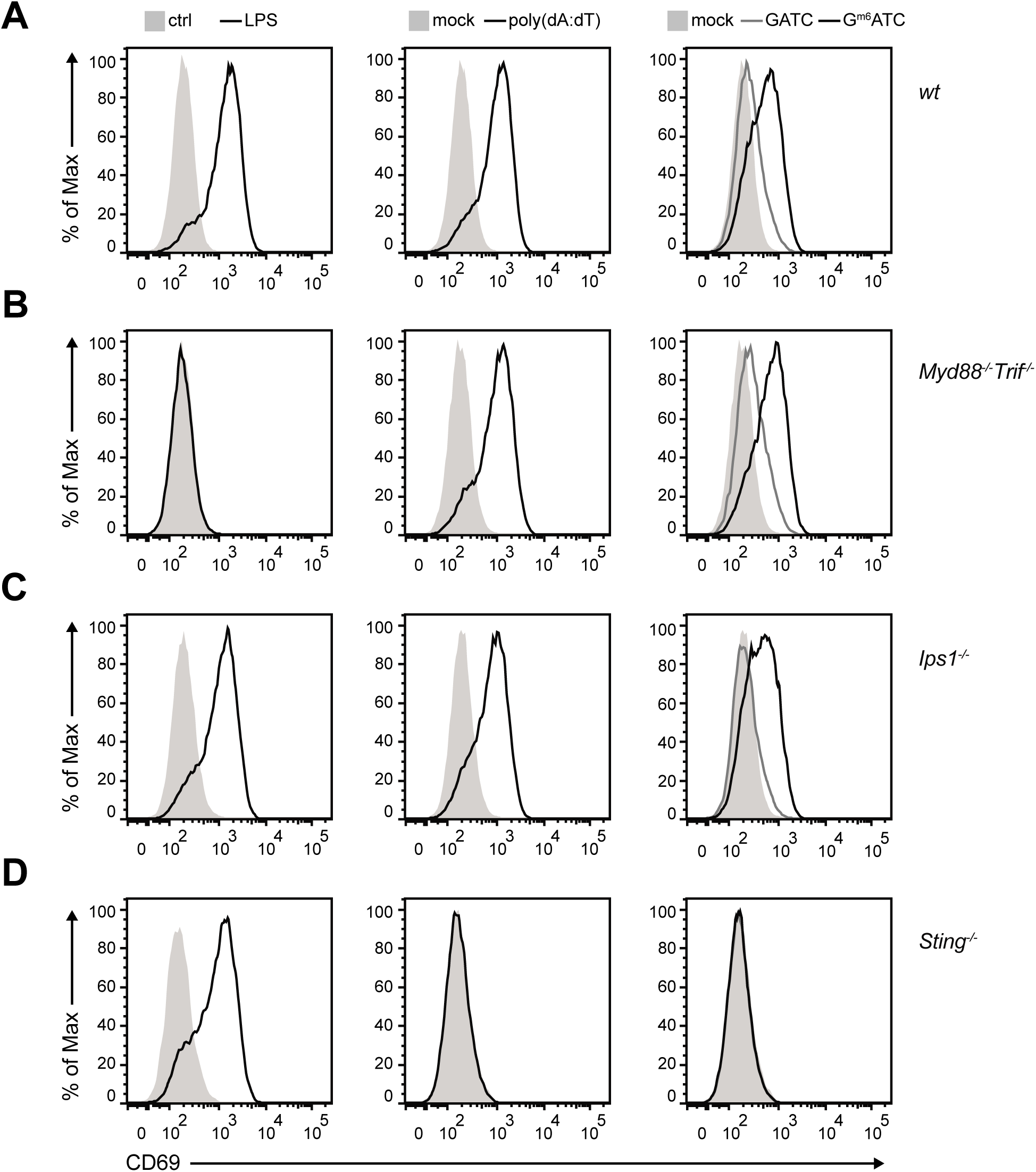
STING is required for macrophage activation by dsDNA irrespective of methylation status. CD69 expression levels determined by flow cytometry of BMMs from (A) *wt*, (B) *Myd88*^-/-^*Trif*^-/-^, (C) *Ips1*^-/-^ or (D) *Sting*^-/-^ mice activated for 6 h with 1 μg/ml LPS, or left untreated (Ctrl; *left panels*). Alternatively, BMMs were transfected with poly(dA:dT) or mock-transfected (*middle panel*), or were transfected with GATC and G^m6^ATC DNA, respectively (*right panel*). Data are representative of two independently performed experiments.

STING was identified as a key adaptor molecule of cytosolic DNA sensing (27). In line with this, transfection of *Sting*^-/-^ BMMs with dsDNA did not result in expression of CD69 protein upon transfection (Fig. 2D). The lack of recognition occurred independently of the m6A modification (Fig. 2D). Thus, STING is required to recognize cytosolic dsDNA, and this recognition is permissive to epigenetic modifications within the DNA.

### Enhanced BMM-activation by m6A methylated DNA is sequence specific

We then interrogated whether the increased immunogenicity of G^m6^ATC DNA was a general feature of m6A methylated DNA. In fact, in addition to the GATC sequence-specific Dam Methyltransferase (MTse), a number of other m6A DNA MTses have been described (16, 18, 34). For instance, *Thermoplasmata* express a m6A MTse that recognizes CATG sequences (16). Another m6A MTse found in *Helicobacter pylori* recognizes adenine within GTAC motifs (35). To determine whether m6A methylations within these motifs also increased the immunogenicity of DNA, we generated dsDNA with the identical 34 bp core sequence, but with the GATC motifs exchanged to m6A-methylated or unmethylated CATG and GTAC motifs (Table 2). Similar to the GATC containing DNA, C^m6^ATG and GT^m6^AC DNA displayed a reduced Tm compared to the respective unmethylated dsDNA (Table 2), indicating that m6A methylation also affects the strength of dsDNA bonds in these sequences.

Comparable to G^m6^ATC DNA, transfecting BMMs with DNA containing GT^m6^AC also induced higher CD69 expression levels than its unmethylated counterpart (Fig. 3A). However, this was not the case for C^m6^ATG DNA. Transfecting BMMs with DNA containing C^m6^ATG resulted in lower CD69 expression than transfection with the unmethylated DNA (Fig. 3A). Furthermore, whereas G^m6^ATC and GT^m6^AC were also superior in increasing *Ifnβ, iNos and Cxcl10* transcript levels compared to the respective unmethylated DNA, C^m6^ATG-containing DNA rather hampered the induction of these key inflammatory genes (Fig. 3B-D). Thus, the observed enhanced immunogenicity of m6A methylation in DNA sequences is sequence-specific.

**Figure 3.**
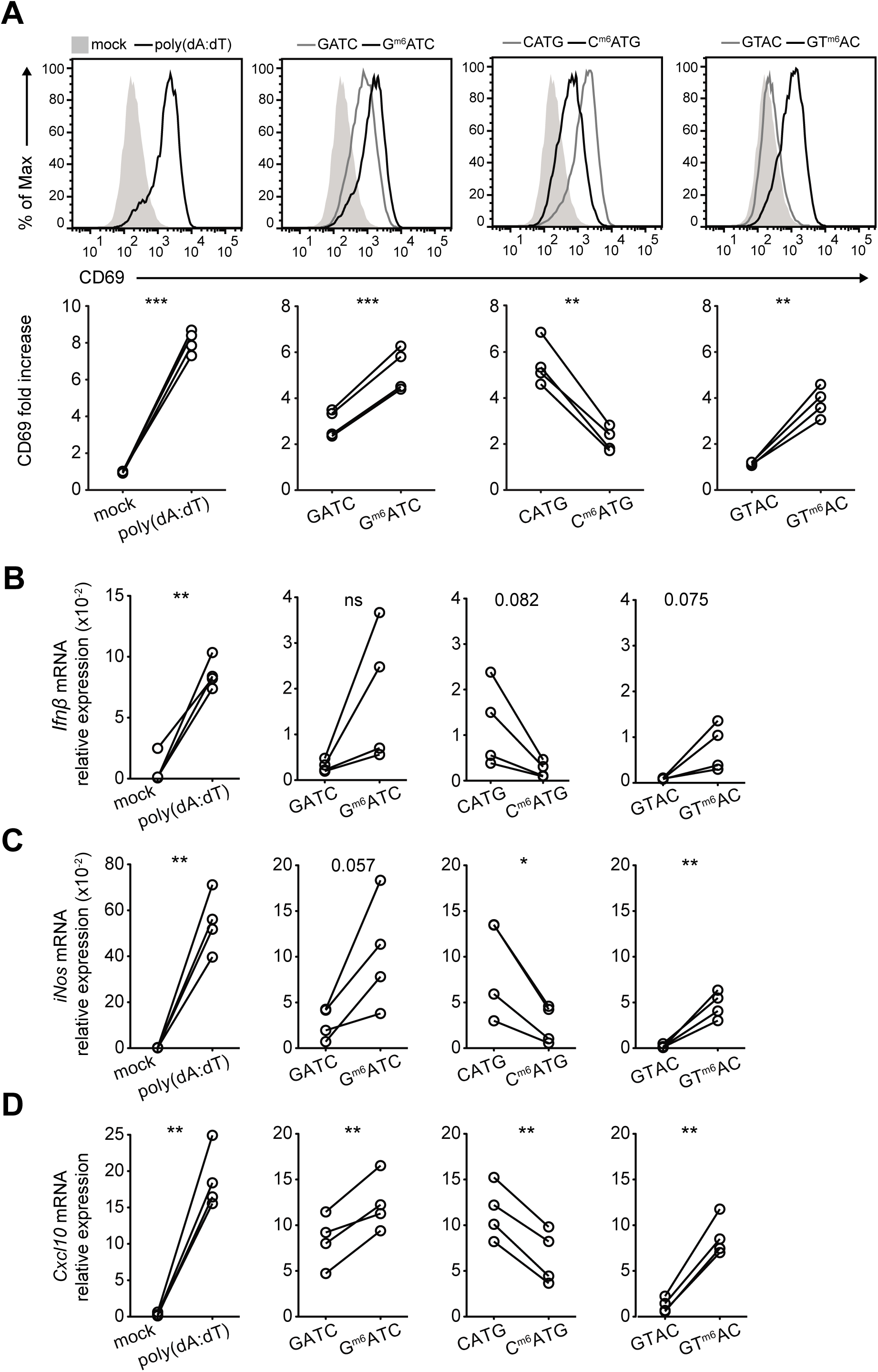
BMMs recognize m6A-methylated dsDNA in a sequence-dependent manner. **(A)** BMMs were mock transfected or transfected for 6 h with poly(dA:dT), (*left panel*), with GATC or G^m6^ATC DNA (*second panel*), CATG or C^m6^ATG (*third panel*), or GTAC or GT m6AC DNA (*right panel*). For sequences see Table 2. Top row: Representative histograms of CD69 expression measured by flow cytometry. Bottom row: Compiled data from BMM cultures of four mice from two independently performed experiments. (**B-D**) mRNA levels of *Ifnβ* (**B**) *iNos* (**C**), and *Cxcl10* (**D**) in BMMs after 6 h stimulation with indicated reagents, normalized to the expression of *L32*. Paired Student’s *t*-test. (* *p* < 0.05, * * *p* < 0.01, * * * *p* < 0.001. ns = not significant).

### Sequence-specific recognition of m6A methylated DNA is conserved in human macrophages

To determine whether the observed differences in sequence-specific immunogenicity was also found in humans, we generated M-CSF derived macrophages from peripheral blood derived monocytes and compared the gene expression levels of effector molecules upon DNA transfection. Comparable to murine macrophages, transfecting human macrophages with G^m6^ATC-containing DNA resulted in higher induction of *CXCL10* mRNA compared to unmethylated DNA (Fig. 4A). The increased immunogenicity of DNA was also conserved for GT^m6^AC DNA (Fig. 4A). In contrast, transfecting macrophages with C^m6^ATG DNA again lowered the induction of *CXCL10* mRNA (Fig. 4A).

**Figure 4:**
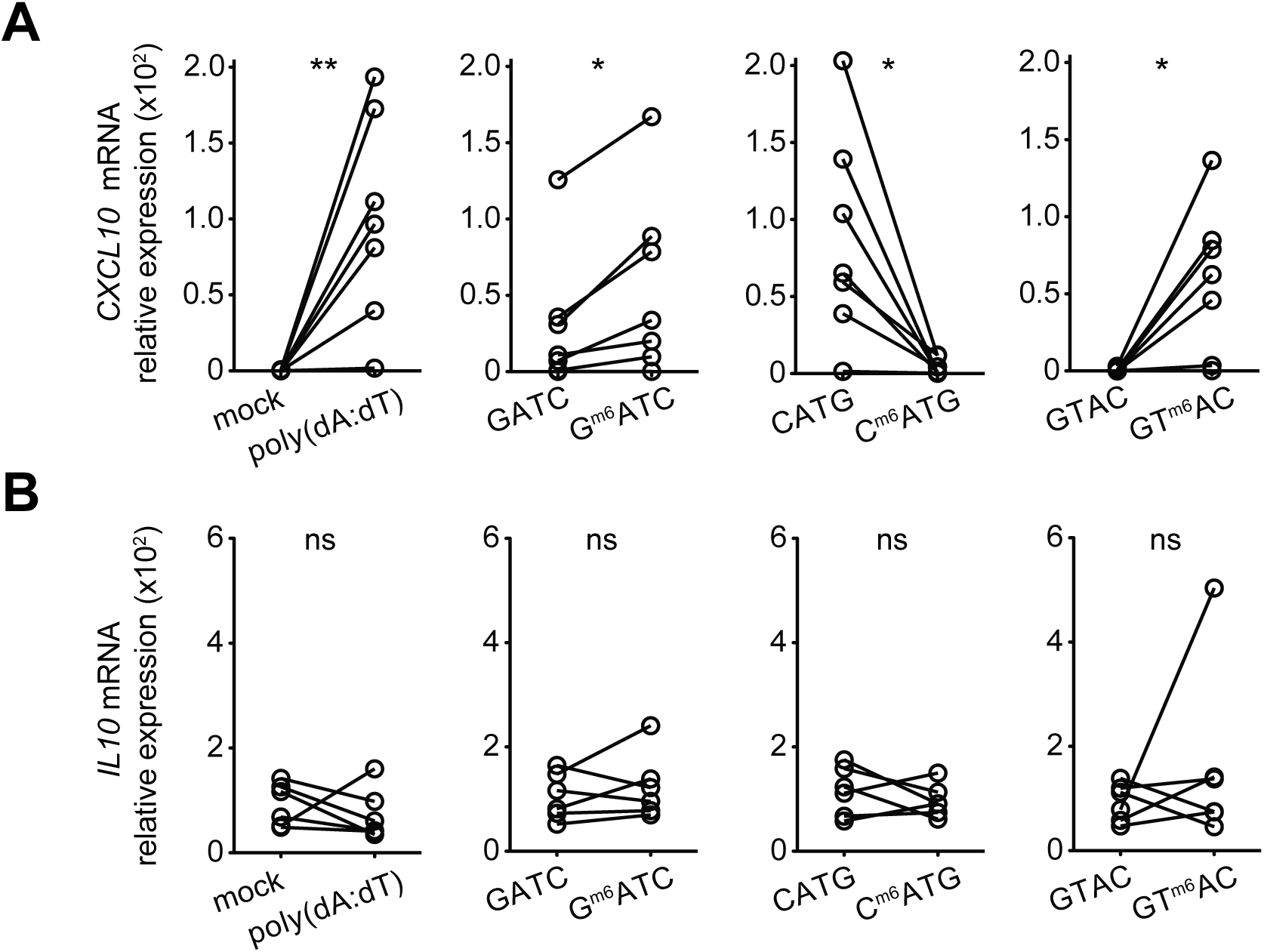
Sequence-specific recognition of m6A-methylated dsDNA is conserved in human macrophages. **(A**, **B)** M-CSF induced macrophages from human peripheral blood derived monocytes were transfected with poly(dA:dT) (*left panel*), GATC or G^m6^ATC DNA (*second panel*), CATG or C^m6^ATG (*third panel*), or GTAC or GT ^m6^AC DNA (*right panel*). mRNA levels of *CXCL10* (**A**) and *IL10* (**B**) were measured and normalized to the expression of *18S*. n = 7 independent donors, measured in four independently performed experiments. Paired Student’s *t*-test. (* *p* < 0.05, * * *p* < 0.01. ns = not significant).

Because the C^m6^ATG sequence in transfected DNA blocked the induction of pro-inflammatory molecules in macrophages, we investigated whether this sequence instead induced the expression of a prototypic anti-inflammatory cytokine, IL-10. However, we did not detected increased *IL10* mRNA levels with any of the m6A methylated DNA sequences when compared to mock-transfected cells (Fig. 4B). In conclusion, the sequence-specific immunogenicity by m6A-methylated DNA motifs is conserved between mouse and human.

## Discussion

Recognition of intracellular dsDNA is an important process that can occur during microbial infection and after cell damage (3). Whereas length and structure modulate the immunogenicity of DNA, we show here that this is also true for m6A-methylation. This increased antigen recognition is observed at all different tested time points and doses. Immune recognition of m6A-methylated DNA is identical to unmethylated DNA in that it is independent of MyD88/TRIF and IPS-1 signaling but requires STING. Which molecule recognizes the sequence and how the m6A methylation influences the immunogenicity is yet to be determined. Interestingly, in *E. coli*, m6A-methylation was shown to affect the oligonucleotide structure and - as a consequence - the binding to the DNA binding protein IHF (36). Here, we observed different Tm in the presence or absence of m6A-methylation in the dsDNA, indicative for alterations in the secondary structure of DNA. This alteration could potentiate the binding affinity of DNA to its cytosolic receptor. Although less likely, m6A-methylation could also lead to limited recognition by cytosolic nucleases and thus support a prolonged exposure to DNA sensors, or enhanced transfection efficiency of DNA. It is tempting to speculate that C^m6^ATG motifs differ in structure from G^m6^ATC and GT^m6^AC motifs, which interferes with recognition of DNA sensors and thus dampens the immunogenicity of dsDNA.

In conclusion, our study identifies a new role for m6A-DNA methylation in regulating innate immune responses to cytosolic DNA. Whether the observed sequence-specific recognition of m6A-methylated DNA is a specific feature of synthetic DNA or stems from different immune responses to various bacterial strains is yet to be determined. Overall, our findings may help to increase the immunogenicity of DNA vaccines and could potentially pave the way to unravel novel mechanisms of pathogen recognition and evasion in innate immune cells.

## Acknowledgements

We thank A. Popovski for technical help, and S. Naik for providing murine Flt3L to generate BM-derived dendritic cells, and J. Freen-van Heeren for critical reading of the manuscript. The 2.4G2 antibody was kindly provided by Louis Boon (Bioceros, Utrecht).

